# Multidimensionality of plant defenses and herbivore niches: implications for eco-evolutionary dynamics

**DOI:** 10.1101/070250

**Authors:** Nicolas Loeuille, Céline Hauzy

## Abstract

Plant defenses are very diverse and often involve contrasted costs and benefits. Quantitative defenses, whose protective effect is dependent on the dose, are effective against a wide range of herbivores, but often divert energy from growth and reproduction. Qualitative defenses often have little allocation costs. However, while deterrent to some herbivores, they often incur costs through other interactions within the community (eg, decrease in pollination or attraction of other enemies). In the present work, we model the evolutionary dynamics of these two types of defenses, as well and the evolutionary dynamics of the herbivore niche. We assess the effects of such evolutionary dynamics for the maintenance of diversity within the plant-herbivore system, and for the functioning of such systems under various levels of resource availability. We show that the two types of defenses have different implications. Evolution of quantitative defenses often helps to maintain or even increase diversity, while evolution of qualitative defenses most often has a detrimental effect on species coexistence. From a functional point of view, increased resource availability selects for higher levels of quantitative defenses, which reduces top-down controls exerted by herbivores. Resource availability does not affect qualitative defenses, nor the evolution of the herbivore niche. The growing evidence that plant defenses are diverse in types, benefits and costs has large implications not only for the evolution of these traits, but also for their impacts on community diversity and ecosystem functioning.

## Introduction

Understanding the evolution of plant defenses is of great importance for ecology and its applications. Because plants serve as the energetic basis of most ecosystems, defenses, by modifying the strength of top-down controls (Chase et al., 2000; Loeuille and Loreau, 2004; Schmitz et al., 2000) may alter the availability of this energy for higher trophic levels (Dickman et al., 2008). Plant defenses also play a critical role in the community composition, not only of herbivores (Becerra, 2007; Kessler et al., 2004; Robinson et al., 2012; van Zandt and Agrawal, 2004; Whitham et al., 2003), but also of higher trophic levels (Halitschke et al., 2008; Poelman et al., 2008; Xiao et al., 2012) and of pollinator assemblages (Adler et al., 2006, 2012; Herrera et al., 2002).

While many works study the coevolution of plants and enemies (Agrawal and Fishbein, 2008; Bergelson et al., 2001; Carroll et al., 2005; Cornell and Hawkins, 2003; Loeuille et al., 2002; Rausher, 2001, 1996), current ecological theory linking the evolution of plant defenses to community structure in general is scarce. Also, from an evolutionary point of view, the fitness components incorporated in such studies are often too simplistic to account for community aspects efficiently. Particularly, most studies focus on the evolution of plant defenses assuming allocation costs (de Mazancourt et al., 2001; Loeuille and Loreau, 2004; Loeuille et al., 2002), proposing that additional defenses divert energy from growth and reproduction (Coley, 1986; Herms and Mattson, 1992; Züst et al., 2011). Such defenses have far reaching implications for ecosystem functioning because they largely decrease the availability of energy for higher trophic levels in two ways. First, by protecting plant biomass, these defenses constrain the proportion of productivity transmitted up the food chains. Second, these defenses reduce the productivity, because of direct allocation costs.

When food chain length is constrained by energy availability (Dickman et al., 2008; Oksanen et al., 1981; Pimm and Lawton, 1977; Wollrab et al., 2012), such costs ultimately modify the structure of ecological networks.

While allocation costs have been widely observed for such quantitative defenses (Müller-Schärer et al., 2004; Strauss et al., 2002), whose efficiency is typically dependent on the dose produced by the plant (for chemical defenses) or for the quantity of protective structures (eg, hair, spines), several studies failed to detect such allocation costs (Häring et al., 2008; Koricheva et al., 2004). A possibility is that allocation costs exist but were not properly detected, these defenses may also be constrained by alternative costs, for instance through other ecological interactions (ecological costs: Müller-Schärer et al., 2004; Strauss et al., 2002). A higher investment in such defenses can be efficient against some enemies, but incurs costs by attracting other enemies or by rendering the plant less attractive to mutualists (e.g., Adler *et al.* 2012; Xiao *et al.* 2012). Ecological costs may be particularly suitable for qualitative defenses (Müller-Schärer et al., 2004; Strauss et al., 2002), for which the presence of the compound rather than its concentration matters for herbivore deterrence. For instance, some volatile compounds seem to be very variable and efficient only against a given herbivore specialist (Becerra, 2003). Many closely related volatile organic compounds exist (Courtois, 2010), involving similar chemical structures and enzymatic pathways. Switching from one to another likely does not incur a large cost in terms of growth or reproduction. While defenses with ecological costs do not have the direct energetic implications of defenses based on allocation costs, their variations largely impact relative interaction strengths within the community. They can also play a crucial role in the diversification of herbivore and plant clades (Becerra, 2007, 2003).

In the present article, we aim at understanding the interplay of these two defense types as well as their implications for the evolution of the herbivore. The model we develop contains a qualitative defense that is intimately linked to the herbivore niche, thereby allowing for ecological costs (in the sense that efficiency against one herbivore will come at a cost given another herbivore), and a quantitative defense that reduces any herbivore pressure, whose allocation cost entails a decrease in the plant biomass production. We investigate how evolution of these two defense types and of the herbivore, affect the functioning and structure of the community. More specifically, we ask:

1. Whether the evolution of each defense type alter the persistence of the herbivore in different ways. According to observations detailed earlier, we hypothesize that qualitative defenses may allow the herbivore persistence while quantitative defenses can only be detrimental to it by reducing energetic availability.
2. Whether the evolution of each defense types produces diversification in the plant compartment (ie, the coexistence of different defensive strategies).
3. How the evolution of each defense type affects the functioning of the system, that is the distribution of biomasses among the two trophic levels and its changes with resource availability. We hypothesize that investment in quantitative defenses, by reducing overall vulnerability, will lower top-down controls therefore allowing plant biomass increase (and low response of herbivore biomass).

### Ecological model

We model the dynamics of plant and herbivore biomass (*P* and *H* respectively) within an isolated ecosystem. In the absence of herbivores, we assume that the plant biomass is constrained by a limiting factor (e.g., energy, limiting nutrient, space) and reaches an equilibrium constrained by *K* (carrying capacity).

The intrinsic growth rate of plants is noted *r*. Herbivores consume plants at a rate *β* and converts a proportion *f* of consumed plant biomass into herbivore biomass. We assume that plant growth is limited by direct competition among plants (*α/K: per capita* competition rate). Herbivore mortality rate *m* is constant.

Accounting for these hypotheses, we model the variations in plant and herbivore biomasses over time through a simple Lotka-Volterra system:

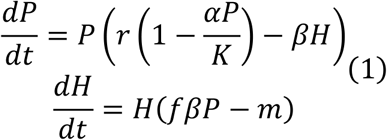

For more details on parameters and variables, see Table 1.

**Table 1:** Notation, name and dimension of variables and parameters

|  | Name | Definition domain | Dimension |
| --- | --- | --- | --- |
| *Variables* |  |  |  |
| $P$ | Plant Biomass | $[0, +\infty[$ | $\text{kg.m}^{-2}$ |
| $H$ | Herbivore Biomass | $[0, +\infty[$ | $\text{kg.m}^{-2}$ |
| $x$ | Plant qualitative defenses | $] -\infty, +\infty[$ | dimensionless |
| $y$ | Plant quantitative defenses | $] -\infty, +\infty[$ | dimensionless |
| $p$ | Herbivore preference (preferred qualitative defenses) | $] -\infty, +\infty[$ | dimensionless |
| $g$ | Degree of generalism of the herbivore | $] 0, +\infty[$ | dimensionless |
| *Functions* |  |  |  |
| $K$ | Carrying capacity | | $\text{kg.m}^{-2}$ |
| $\beta$ | Per capita consumption rate | | $\text{m}^2.\text{kg}^{-1}.\text{time}^{-1}$ |
| $\alpha$ | Trait dependent competition scaling | | dimensionless |
| *Parameters* |  |  |  |
| $K_0$ | Basal carrying capacity of plant | $] 0, +\infty[$ | $\text{kg.m}^{-2}$ |
| $f$ | Conversion efficiency | $[0, +\infty[$ | Dimensionless |
| $m$ | Herbivore *per capita* mortality rate | $[0, +\infty[$ | $\text{time}^{-1}$ |
| $r$ | Maximal plant intrinsic growth rate | $[0, +\infty[$ | $\text{time}^{-1}$ |
| $a$ | Benefits of quantitative defenses in terms of reduced consumption | $[0, +\infty[$ | dimensionless |
| $b$ | Costs of quantitative defenses in terms of reduced competitive ability | $[0, +\infty[$ | dimensionless |
| $\beta_0$ | Basal herbivore consumption rate | $[0, +\infty[$ | $\text{m}^2.\text{kg}^{-1}.\text{time}^{-1}$ |
| $\sigma$ | Variance of the competition kernel | $]0, +\infty[$ | dimensionless |

### Traits and trade-offs

Because plants are consumed by herbivores, herbivores exert a selective pressure on plant defensive traits. The traits of herbivores, whose reproduction and growth depend on the plants they consume, are similarly likely to evolve in response to plant defenses. Hence, the consumption rate of herbivores *β* is shaped by both plant and herbivore traits. We consider that plants are characterized by two defense traits noted *x* and *y*. The consumption strategy of herbivores is characterized by two traits *p* and *g*. Hence, the consumption rate of herbivores *β* is a function of these four traits:

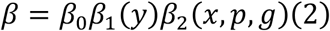

where *β*_0_ is the basal rate of consumption.

Trait *y* represents a quantitative defense that has an allocative cost (Müller-Schärer et al., 2004). The efficiency of trait *y* depends on its amount within each plant. We assume it decreases the herbivore consumption rate:

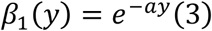

We suppose that allocative costs affect the plant competitive ability (Agrawal et al., 2012):

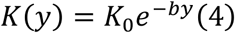

Combining (3) and (4) allows flexible trade-off shapes between investment in defenses (*-β*) and *K:* concave (*a>b*), linear *(a=b)* or convex *(a<b)* (Fig 1A).

**Figure 1.**
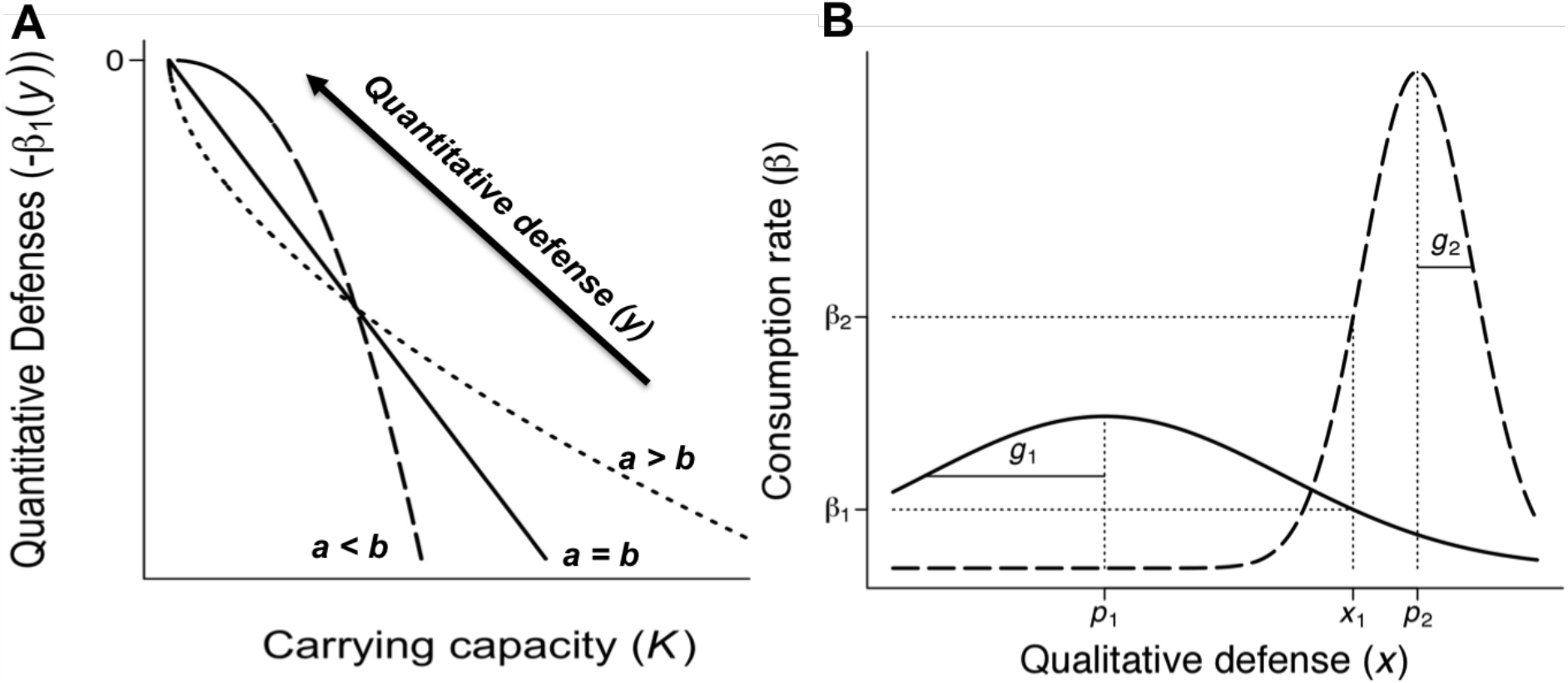
Types of defense and their costs. A. The quantitative defense trait *y* decreases consumption *β* of plants by herbivores and affects competitive ability, lowering the plant carrying capacity *K*. The trade-off can be concave (dashed line, *a<b*), linear (solid line, *a=b*) or convex (dotted line, *a>b*). B. Plant qualitative defense trait *x* of plants defines one dimension of the herbivore niche. Herbivore niche is described by two consumption traits: *p* and *g*. Herbivore preferencep, is the *x* value at which the consumption rate of the herbivore is maximal (trait matching). The generalism of the herbivore, denoted *g*, sets the ability of the herbivore to consume plants a given range of *x* around *p*. The herbivore defined by (*p*_1_,*g*_1_) is a generalist (solid line) whereas the herbivore (*p*_2_,*g*_2_) is a specialist (dashed line). The more generalist the herbivore, the lower is its maximal consumption rate.

Trait *x* represents a qualitative defense. For instance, *x* may be construed as a particular assembly of defensive compounds (*e.g.*, a given chemical bouquet of volatile organic compounds). Each plant is characterized by one qualitative defense value. This qualitative defense *x* defines one dimension of the ecological niche of herbivores (Fig 1B). Along this niche dimension, we consider that herbivore consumption is described by two traits, *p* the preference of the herbivore for a given chemical bouquet and *g* the degree of generalism (*g*>0). The further the herbivore preference *p* is from plant trait *x*, the lower its consumption rate (ie, qualitative defenses affect herbivore consumption through trait matching rules). Herbivore generalism *g* describes the range of trait *x* that can be efficiently consumed by the herbivore. We assume a trade-off between the generalism *g* and the maximal consumption rate (Craig MacLean et al., 2004), so that the consumption rate is normalized and remains globally constant when *g* varies. Accounting for these constraints, the herbivore niche is (Fig 1B):

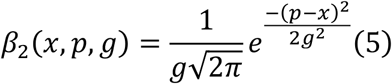

Note that trait *x* does not entail any direct cost. However, changes in *x* may be constrained by ecological costs (ie, by increasing interaction with other herbivores). For instance, if plant trait *x* is between the traits of two herbivores (*p_1_* and *p_2_* respectively), then any variation of *x* will decrease the interaction with one herbivore, but attract the other (eg, as on Fig 1B). As a result of equations 4 and 5, we have two traits for defense: one with allocation costs and no ecological cost (*y*), while the other has only ecological costs and no allocation costs (*x*). We acknowledge that in nature, defense traits are not likely to be as clear cut, and that qualitative defenses may actually involve weak allocation costs or quantitative defense may be counteracted by some herbivores. However, this simplification allows us to fully describe resulting evolutionary dynamics and to highlight the consequences of various cost structures for plant defenses.

We studied two competitive scenarios: (1) α=1; (2) direct competition is enhanced when traits are similar (Brännström et al., 2011; Kisdi, 1999; Loeuille and Loreau, 2005; Yoder and Nuismer, 2010). We modeled the relationship between the direct competition coefficient *α* and plant traits using a Gaussian function. Similarity is defined by the Euclidean distance *D* between plant traits:

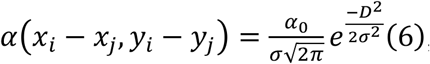

with

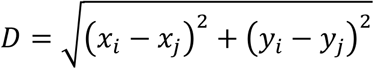

### Evolutionary dynamics

We studied the evolution of plant and herbivore traits using adaptive dynamics methods (Dieckmann and Law, 1996; Geritz et al., 1998). While all traits may coevolve, we here study the evolution of each species and each trait separately, to contrast the implications of the different evolutionary dynamics. Therefore, for each trait, we consider a monomorphic population and determine a fitness of a mutant whose value for the given trait slightly differs.

The relative fitness of a mutant population in a resident population, denoted *W_m_*, depends on both the mutant and resident traits. It is defined as the *per capita* growth rate of a rare mutant population in a resident population at equilibrium (*P*\*, *H*\*). For instance, considering the trait *y*, a mutant plant has a relative fitness:

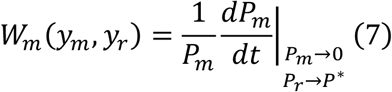

where *y_m_* is the trait of the mutant population while the resident population *P_r_* is assumed to be at equilibrium (ecological dynamics are therefore assumed faster than evolutionary dynamics).

The evolution of a trait is modeled using the canonical equation of the adaptive dynamics, which assumes that the amplitude of mutation effects, *ω*, is small. For trait *y*:

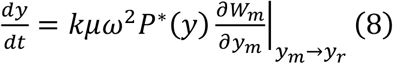

where *µ* is the *per capita* mutation rate, *ω^2^* is the variance of mutation effect, and *k* is a scaling parameter. The selection gradient 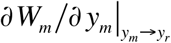 corresponds to the slope of the local adaptive landscape (ie, close to the resident trait) and constrains the direction of evolution. Singular strategies *y*\*, therefore correspond to:

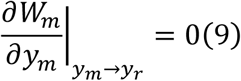

Evolutionary dynamics around the singular strategies can be analyzed by computing the second derivatives of the fitness function (Dieckmann and Law, 1996; Geritz et al., 1998). Singular strategy *y*\*, cannot be invaded by nearby mutants, provided:

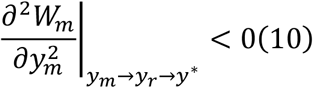

Moreover, singular strategies satisfy the convergence criteria (ie, selection favors mutant closer to the singularity in its vicinity) provided:

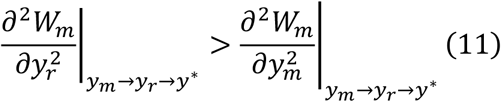

When an evolutionary equilibrium satisfies both the non-invasibility and the convergence criteria, it is called a Convergence Stable Strategy or CSS (Eshel, 1983). When an evolutionary equilibrium satisfies the convergence condition but is invasible, the selection near the equilibrium is disruptive and an evolutionary branching eventually occurs, creating a diversification in the corresponding trait (ie, the coexistence of two or more phenotypes exhibiting different defense levels). Finally, we also encountered singularities that were invasible and non convergent, called repeller.

Note that the framework we use here can be extended to account for variations not in one trait at a time, but of multiple traits simultaneously (eg, Loeuille et al. 2002). It can also be extended to follow the evolution of traits along branches passed the first branching point. Our study can then be thought as the first step of a more complete evolutionary analysis. Our analysis of single traits however allows a complete mathematical analysis of the singularities and associated evolutionary dynamics (detailed in the appendix). More complex coevolutionary scenarios do not allow a tractable analysis of the evolutionary trajectories, as convergence and invasibility criteria cannot be easily extended in such instances (Kisdi 2006).

## Results

We here describe the main results of the analysis. More details, including regarding the formulation of fitness functions, fitness gradients and evolutionary singularities are shown in the supplementary information.

### Ecological dynamics

The model described by the system of equation (1), has a single equilibrium allowing the coexistence of plants and herbivores:

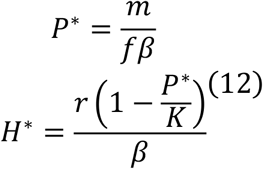

From the Jacobian matrix of (1) estimated at equilibrium (12), it is possible to show that this coexistence equilibrium is stable when it is feasible, i.e. when

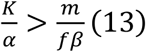

When 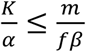, herbivores go extinct and plants reaches 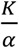.

Effects of enrichment on equilibrium (12) can be studied from derivatives:

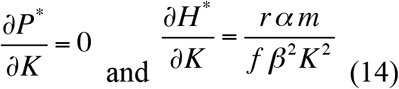

Thus, when considering only ecological dynamics, as plants limiting factor increases (*K* increases), herbivore biomass increases whereas plant biomass remains constant (Fig. 4A), stressing the importance of top-down controls in the ecological model.

### Evolution of quantitative defenses

When the carrying capacity of plants is sufficiently high to maintain herbivores, the consumption of plants by herbivores depends on the level of quantitative defense of plants y. Incorporating trait *y* in equation (13), one gets that herbivore coexist with plants when

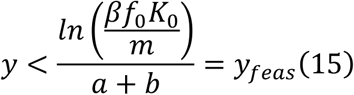

The fitness of a rare plant mutant of trait *y*_m_ in the resident plant population of trait *y*_r_ is then:

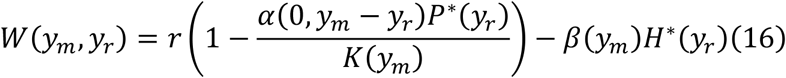

The evolutionary dynamics of the quantitative defense y are described by the canonical equation (8)

The associated singular strategy is:

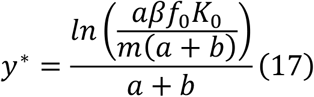

Comparing (17) and (15) shows that the evolutionary singular strategy is always feasible (*y^*^ < y*_feas_). The properties of this evolutionary equilibrium (invasibility and convergence criteria) depend on the competition scenario that is considered. When competition does not depend on trait similarity *(α=1)*, the singular strategy satisfies both the convergence (eq 11) and the non-invasibility (eq 10) criteria, being therefore a Continuously Stable Strategy, or CSS (Marrow et al., 1996). Quantitative defense levels then evolve to reach *y^*^* at which point the evolutionary dynamics stabilize. Note that the selected amount of quantitative defenses increases with energetic parameters of the plant population (eg, *K_0_*) and with herbivore consumption pressures *(β f_0_)*.

By contrast, when the direct competition between plants increases with trait similarity (eq 6), the evolutionary outcome depends on the following condition:

– If 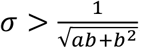, the singular strategy *y^*^* remains a CSS (Fig.2A,B).
– If 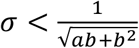, the singular strategy *y^*^*, while still convergent becomes invasible. In such instances, disruptive selection yields successive evolutionary branchings leading to the coexistence of a diversity of quantitative defense strategies, ie the coexistence of plant phenotypes exhibiting contrasted levels of quantitative defenses (Fig.2C,D). Note that in such instances, values of the trait y can become larger that the limit value *y_feas_* (eq 15). Eq 5 is indeed computed from the one plant-one herbivore system (eq 12), while on Fig 2C, the herbivore consumes a set of plants exhibiting various defense traits y, including one abundant plant species that is palatable (the y of the lower branch allows a feasible system).

**Figure 2.**
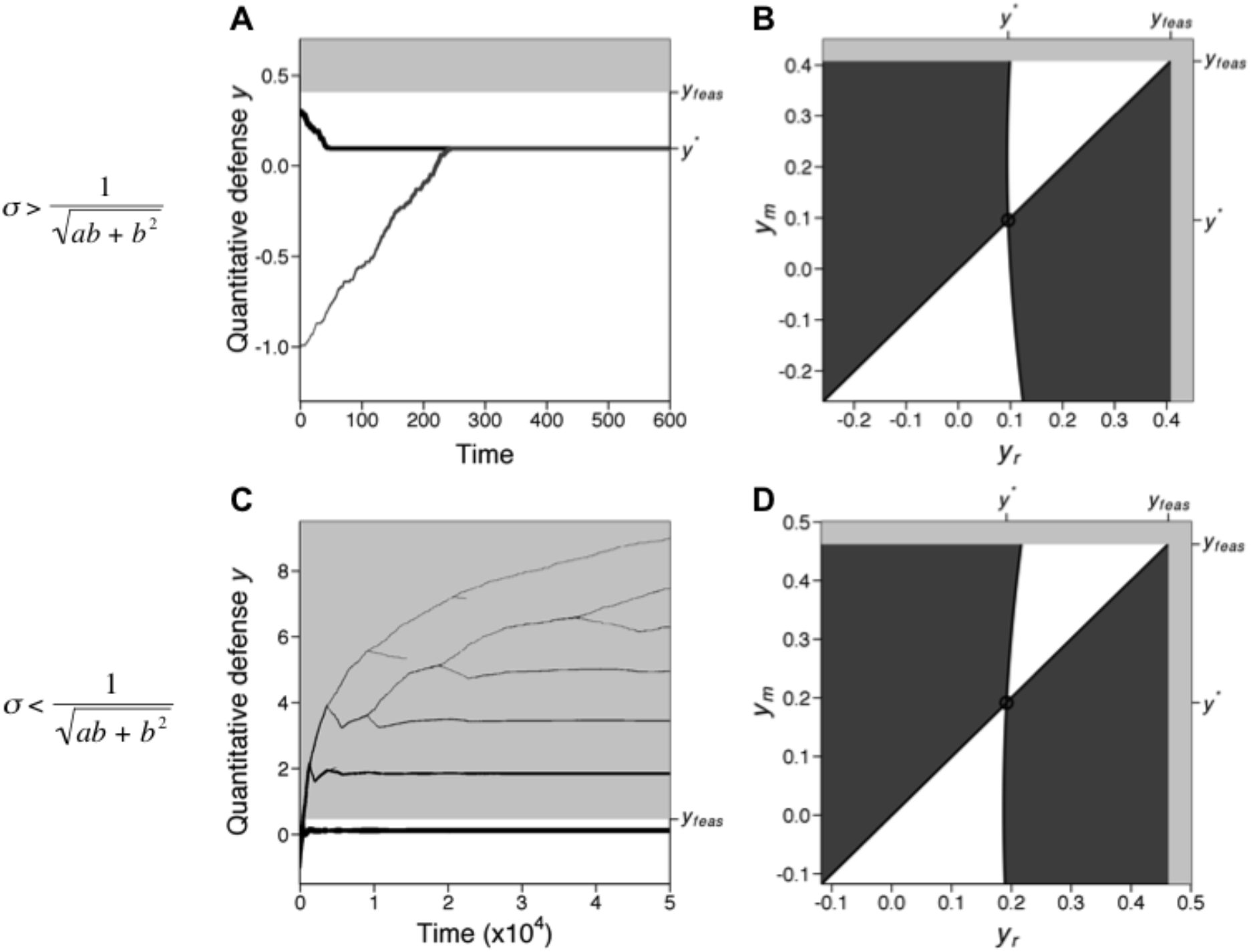
Evolution of quantitative defenses *y* assuming that competition increases with trait similarity. The herbivore, feeding on one plant, maintains a positive biomass *H*\* if the quantitative defense *y* is below *y*_feas_ (*H*\*<0, light grey background; *H*\*>0 white background). When *σ* is high (A, B), trait difference has small effects on the direct competition, the quantitative defense *y* converges to the evolutionary equilibrium *y\** which is a CSS. When *σ* is low (C, D), similar morphs compete very strongly, yielding disruptive selection and successive evolutionary branchings. On A and C, the thickness of lines is proportional to plant biomass. (B, D) Pairwise Invasibility Plots show the sign (+: dark grey area; white area) of mutant fitness as a function of the trait of the resident *y*_r_ and of the mutant *y*_m_. Parameter values (A, B, C and D): *r*=1, *K*_0_=10, *α*_0_=1, *σ*=0.4, *β*_0_=1, *f*=0.1, *m*=0.5, *b*=1. (A,B): *a*=0.7. (C,D): *a*=0.5

Variations in biomasses *P\** and *H\** and in trait *y\** with plant limiting factor can be studied by differentiating with respect of *K*_0_ (see appendix). Contrary to the pattern observed for the purely ecological model, when the evolution of the quantitative defense *y* leads to a CSS, the plant biomass *P*\*, herbivore biomass *H*\* and the level of defense *y*\* at the evolutionary equilibrium all increase with *K*_0_ (Fig. 4B). Evolution of quantitative defenses therefore allows the plants to reduce top down controls exerted by the herbivore.

### Evolution of qualitative defenses

Now fixing quantitative defense level *y*, we analyze the evolution of qualitative defenses. Incorporating *x* in the feasibility condition (13), coexistence is possible if:

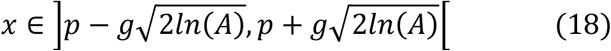

(*A*>1). When direct competition between plants is independent on *x (α=1)*,

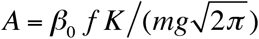. when direct competition between plants depend on plants similarity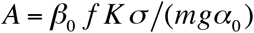.

The only possible singular strategy is ^*x* = P*^ (independent of the competitive scenario). Convergence and non-invasibility criteria are always violated; making this singular strategy a repeller (Geritz et al., 1998).Thus, evolutionary dynamics always move away from herbivore preference *p*. Such an outcome is intuitive. As we assume no direct costs of qualitative defenses *x*, they may only be counterselected when they increase consumption by other herbivores. As our model here just considers one herbivore, plant evolution is continuous and directional. Eventually, the evolution of the qualitative defense leads to herbivores extinction (evolutionary murder *sensu Dercole et al., 2006)*, when *x* reaches the feasibility boundaries (eq 18). It is possible to understand how resource availability affects the ecological and evolutionary states, by differentiating equilibrium biomasses and trait with respect to *K*. Higher levels of resources increase herbivore biomass while plant biomass and plant qualitative defenses *x\** remain unaffected (see appendix & Fig. 4C).

### Evolution of herbivore preference

When the carrying capacity of plants is sufficiently high to maintain herbivores, the consumption of plants by herbivores is constrained by the difference *p-x*. Herbivore biomass is strictly positive if 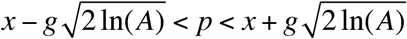 where *A*>1 and 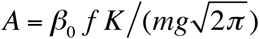.

Only one evolutionary equilibrium then exists, *^p* = x^*, which is always convergent and cannot be invaded (CSS). Evolution eventually leads to this value. Thus, herbivore preference *p* increases or decreases depending on its initial position with respect to *x* until herbivore preference matches plant qualitative defenses *x*. As for plants, such simple dynamics would be altered in more complex communities. An herbivore consuming several plants differing in their trait *x* would face a trade-off between the consumption of one plant and the other.

Higher resource availability leads to an increase of herbivore biomass while plant biomass and herbivore preference *p\** are unaffected (see appendix & Fig. 4D).

### Evolution of herbivore generalism

Equilibrium value of herbivore biomass as defined by equation (12) can be defined as a function of trait *g*, and that this function reaches a peak at |p-x|. This peak is positive (ie, herbivore population can be positive), only if ^*|p-x| <B*^ where 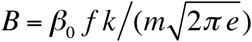.

Therefore, *g* is constrained to an interval 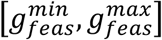 that allows both plant and herbivore populations to be positive.

The singular strategy associated with herbivore specialization is ^*g* = |p - x|*^. This singularity is by definition feasible (see the argument above). This equilibrium satisfies non-invasibility and convergence criteria and is thus a CSS (Fig.3B). Evolution of herbivore generalism *g* therefore converges toward *g\** (Fig.3A). Selection acts to match the degree of the generalism of the herbivore with the difference that exists between its preference and the trait of the available plant population. Differentiating with respect of *K*, it may be shown that any increase in *K* leads to an increase in herbivore biomass while plant biomass and herbivore generalism *g\** are not affected by resource availability (appendix & Fig. 4E).

**Figure 3:**
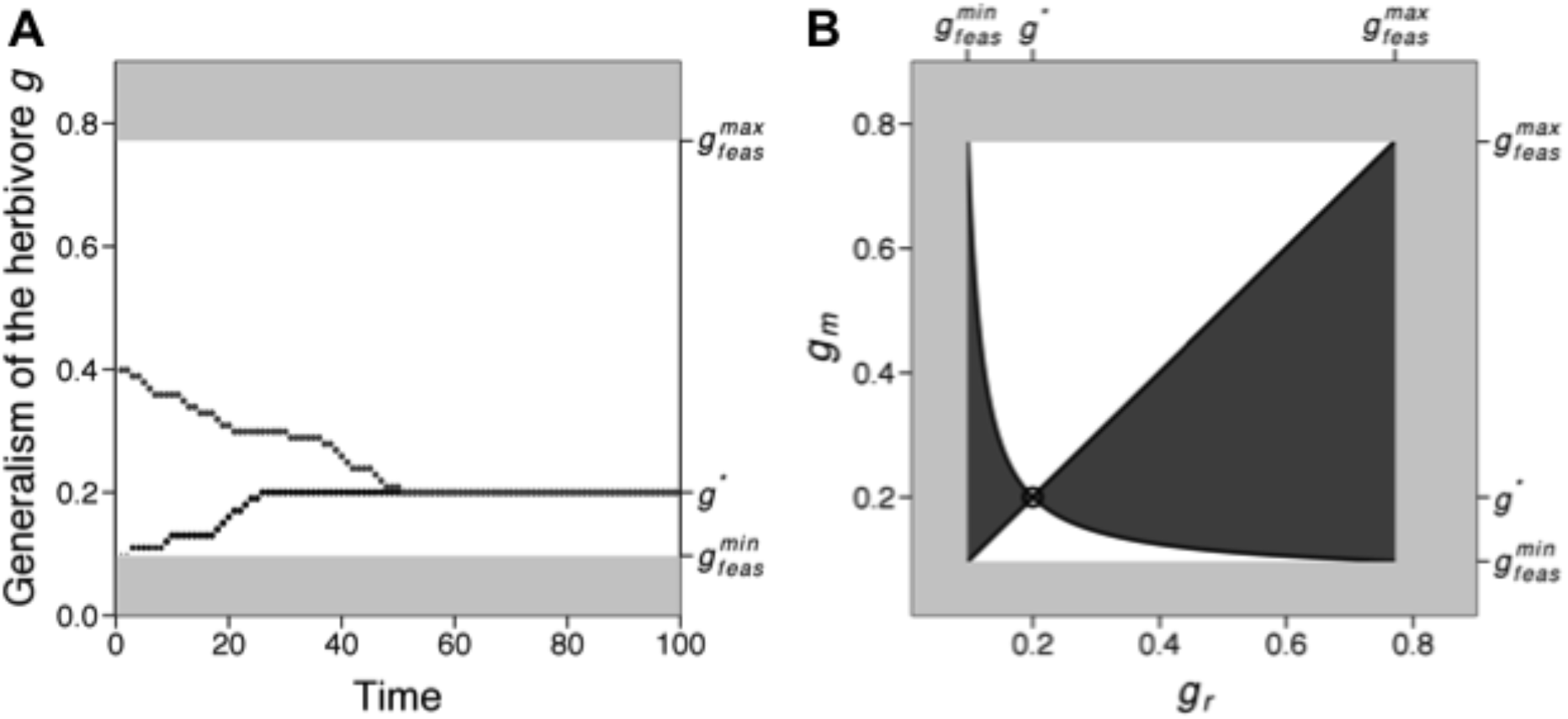
Evolution of herbivore generalism *g*. The herbivore maintains a positive biomass *H*\* if its generalism *g* is between 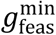 and 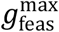 (*H*\*<0, light grey area; *H*\*>0 white area). Generalism converges to an evolutionary equilibrium *g* = |p — x|* that is a CSS. (A) Two examples of evolutionary dynamics for two initial values of *g* (*g*_0_=0.1; *g*_0_=0.4). (B) Pairwise Invasibility Plots near represent the sign (+: dark grey area; -: white area) of mutant fitness as a function of the trait of the resident *g*_r_ and of the mutant *g*_m_. Parameter values (A, B): *r*=1, *K*=10, *α*_0_=1, *σ*=0.4, *β*_0_=1, *f*=0.1, *m*=0.5, *p*=0.3, *x*=0.5.

**Figure 4.**
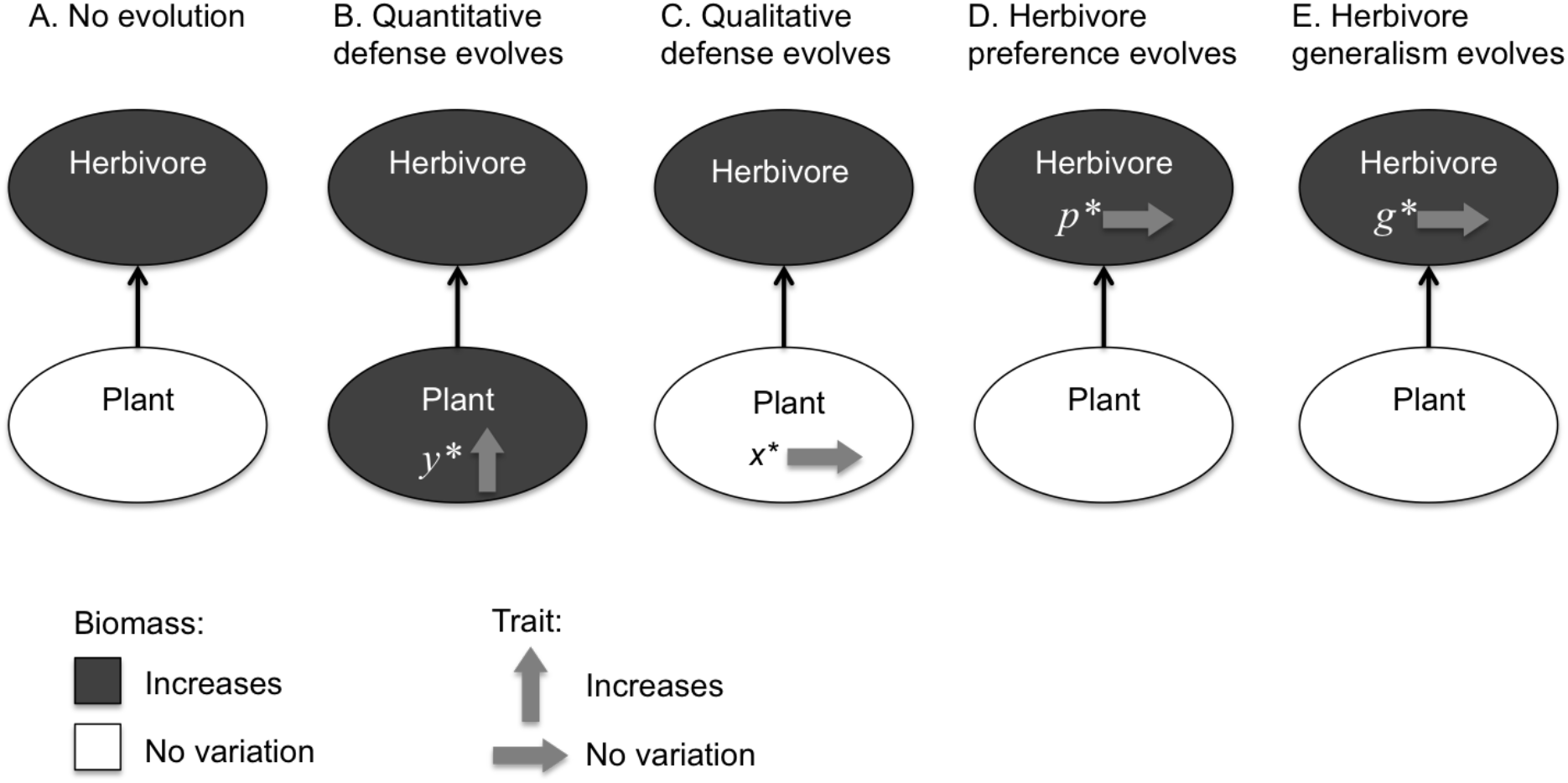
Effects of increases in resource availability, depending on the eco-evolutionary scenario. Without evolution, enrichment has a positive effect on the density of herbivores (A). This pattern remains when the herbivore evolves (D, E) or when qualitative defenses evolve (C). Quantitative defenses (B) are increased when resource levels are higher, allowing for an increase in plant biomass.

## Discussion

The aim of the present work is to understand how the evolution of various types of plant defenses and of herbivore consumption strategies alters the structure and the functioning of plant-herbivore systems. The two types of defenses we consider have been proposed based on reviews of many different empirical systems (Müller-Schärer et al., 2004; Strauss et al., 2002) that distinguish quantitative defenses (efficient against all herbivores, but having allocative costs that reduce growth or productivity) and qualitative defenses (whose costs are not allocative, but happens through the modifications of other interactions). Most theoretical works on plant defenses focus on the former type (de Mazancourt et al., 2001; Levin et al., 1990; Loeuille and Loreau, 2004; Loeuille et al., 2002; Loreau and Mazancourt, 1999), while the evolution of qualitative defenses has received far less attention (but see Loeuille and Leibold, 2008).

Concerning the structure of the community, evolution of quantitative defenses tends to increase the complexity of the system. First, contrary to our prediction, coexistence of the plant-herbivore system is warranted at the evolutionary equilibrium. Evolution of quantitative defenses indeed decreases the herbivore population. At some point, herbivore population becomes too low and selection of higher levels of defense incurs too much intrinsic costs for little benefits. Evolution then stops, but the herbivore persist (through at smaller biomass). Next to maintaining the different trophic levels, the evolution of quantitative defenses also increases the plant phenotypic diversity, when disruptive selection allows the coexistence of plant phenotypes that have contrasted levels of defenses. Such a diversification within the plant compartment however requires that plant competition is partly linked to trait similarity. These results are consistent with other models that predict branching in defense strategies (Costa et al., 2016; Ito and Ikegami, 2006), but also, from an empirical point of view, with the widespread coexistence of contrasted investment in defenses within natural ecosystems (Züst et al., 2012).

Evolution of qualitative defenses, on the contrary, leads the system to simpler structures. Our results suggest that, *per se*, the evolution of such defenses should lead to strategies that ever diverge from the herbivore preference. Because evolution away from the herbivore does not involve costs in itself, evolution eventually allows the existence of plants that will be too little consumed to compensate the herbivore intrinsic mortality rate. Evolution of plant then kills the herbivore (evolutionary murder *sensu Dercole et al., 2006)* thereby constraining the maintenance of diversity within the community. Also, we note that, in the case of qualitative defenses, diversification of defense strategies is never observed, even when similar plants compete more strongly. We therefore suggest that, intrinsically (ie, under our assumption of a simple one plant-one predator community), evolution of qualitative defenses may limit diversity (both in terms of species coexistence and in terms of phenotypic variability) while the evolution of quantitative defenses ultimately favors diversity. Finally, note that, herbivore evolution in response to qualitative defenses, either through variations in its preference or through variations in its generalism, always allows the coexistence of the plant-herbivore community. It however does not lead to the coexistence of various herbivore strategies.

In terms of ecosystem functioning, we uncover the impact of variation in resource availability on the eco-evolutionary dynamics of the plant-herbivore system. In all scenarios, higher levels of resources always increase herbivore biomass. When only ecological dynamics is allowed, the plant biomass remains constant. Such a pattern is expected, as our model formulation allows for strong top-down effects (Hairston et al., 1960; Oksanen and Oksanen, 2000; Oksanen et al., 1981). The evolution of herbivore strategies or of plant qualitative defenses does not alter this pattern. Indeed, evolution of these traits is independent of resource supply, as qualitative defenses do not hinge on allocative costs and herbivore traits define the niche of herbivores based on such qualitative defenses. Evolution of quantitative defenses, on the other hand, is expected to alter the pattern that would be expected when discarding evolution. Higher resource availability relaxes the allocation constraints that affect quantitative defenses. It then allows the production of higher levels of defenses, which in turn decreases the effects of top down controls by modulating the herbivore consumption rate. In such a scenario, plant biomass then increases when more resources are available. Such a weakening of top-down controls due to plant defenses is in good agreement with other theoretical/conceptual works (Armstrong, 1979; Leibold, 1996; Loeuille and Loreau, 2004; Strong, 1992), and has been suggested as an important mechanism for the mitigation of trophic cascades in nature (Borer et al., 2005; Polis et al., 2000). Our results again highlight that considering different types of defenses is especially important to understand the fate of ecosystems undergoing environmental change. Whether plants are defending themselves with qualitative or quantitative defenses eventually leads to contrasted outcomes in terms of ecosystem functioning here. Finally, note that the evolutionary part of these results may also be tested. Along environmental gradients of resources we for instance expect molecules acting as quantitative defenses will systematically increase, while molecules acting as qualitative defenses will remain approximately constant.

While the two types of defenses have contrasted effects on coexistence, one may wonder how their evolutions affect the system stability. As explained in the result section, in the case of our linear model, coexistence of the two species insures that the equilibrium is asymptotically stable. However, it is still possible to assess the return time to the equilibrium (assuming a small disturbance on the equilibrium), through the changes in the eigenvalues of the associated jacobian matrix (eg, Rip & McCann 2011). Earlier works have shown that the resilience of the system will increase when the consumer (here herbivore) death rate increases relative to the its attack rate (Rip & McCann 2011). If one considers the effects of enrichment in the model (figure 4), it is possible to infer how resilience will change depending on the type of defense that evolves. When quantitative defenses evolve, enrichment leads to more defenses (Fig 4B), thus a lower attack rate, so that the system becomes less stable, in line with predictions from the paradox of enrichment (Rosenzweig 1971). Conversely, when qualitative defenses evolve, enrichment does not lead to any change in the evolved defenses, so that attack rates are constant and stability unchanged.

We however stress that the model we use here is deliberately simple as its goal is mostly to contrast eco-evolutionary dynamics linked to various plant-herbivore traits. We expect that two levels of complexity, not considered here, will indeed matter much for most empirical situations. First, it seems likely that most plants do not use quantitative defenses or qualitative defenses, but actually use the two types of defenses simultaneously. Also, while the quantitative/qualitative dichotomy is useful as a first approximation, costs and effects are likely to vary in a more continuous fashion so that defenses actually follow a continuum between the two extremes (qualitative/quantitative) used to structure the present work. When considering the coevolution of quantitative and qualitative defenses, we expect strong interactions between their evolutionary dynamics. Consider for instance that the cost of qualitative defenses is to attract another herbivore. A plant having high levels of quantitative defenses would not pay much of such a cost, for it is protected against such alternative herbivores. Now imagine a fast variation in qualitative defenses (as they are involve little direct costs) in response to increase in an herbivore population. Such fast evolutionary dynamics will negatively impact the herbivore population, thereby decreasing the selective pressures for quantitative defenses. We therefore expect that quantitative and qualitative defenses create evolutionary feedbacks on one another, so that the study of their coevolution is especially interesting and an exciting perspective for future works.

A second important simplification lies in the ecological system we use for our analysis. We have considered one single plant and herbivore population, to allow for a more thorough and tractable analysis of the consequences of the evolution of the different traits. An important perspective is to consider the diffuse coevolution of plants and herbivores within diverse communities. Consider for instance the implications of qualitative defenses for diversity. As mentioned at the beginning of this discussion part, the evolution of such defenses ultimately constrains the diversity in our system, the plant eventually “killing” the herbivore through its evolutionary dynamics. We expect this conclusion to differ when a diversity of herbivores is considered.

Consider that, next to the herbivore we modeled in the result part (that has a preference *p_1_)*, we now consider also a second herbivore, whose preference is *p_2_*. Note that, under such conditions, we expect that the most efficient herbivore will win the competition and eventually exclude the other herbivore (R* rule, Tilman, 1982). For the sake of the argument, suppose that ecological and evolutionary dynamics of the plant is however faster than the herbivore dynamics *(e.g.*, because the generation time of herbivores and plants may be vastly different), so that, on a first approximation, we may consider the herbivore population fixed and study the evolution of qualitative defenses *x* in this context. In the one herbivore context, as earlier, selected defenses diverge from the herbivore preference *p_1_* (hence an expected evolutionary murder of this herbivore, figure 5A). The presence of the second herbivore however halts this runaway evolution (figure 5B) by creating a selective force constraining the evolution of qualitative defenses. It thereby allows the first herbivore to remain in the system (at least on this timescale). Similarly, the evolution of the plant due to the first herbivore facilitates the maintenance of the second herbivore (as the plant trait becomes more similar to its preference *p*_2_). Because this evolution actually leads to an equivalent consumption of the plant by the two herbivores, a neutral coexistence is then possible, so that the two herbivores eventually remain in the system. Though the herbivores compete for the plant from an ecological point of view, indirect effects due to the plant evolution from one herbivore to the other are positive, a situation we call “evolutionary facilitation”. Such positive effects due to evolution have already been shown in other contexts. For instance, Abrams and Matsuda (2005) show that adaptation in the prey can facilitate the persistence of its predator. Such indirect interactions between herbivores through plant defenses have been also been suggested in empirical works. Expression of plant defenses following herbivore consumption has been shown to facilitate some other herbivores, while deterring others, so that defenses strongly affect herbivore diversity maintenance (Poelman et al., 2008). The extension of the model we present here, in a more complex network context, would allow a better understanding regarding the role of plant defenses and of herbivore consumption traits in the maintenance of diversity within natural communities. It may also help the management of biological control in an agricultural context (Loeuille et al., 2013).

**Figure 5:**
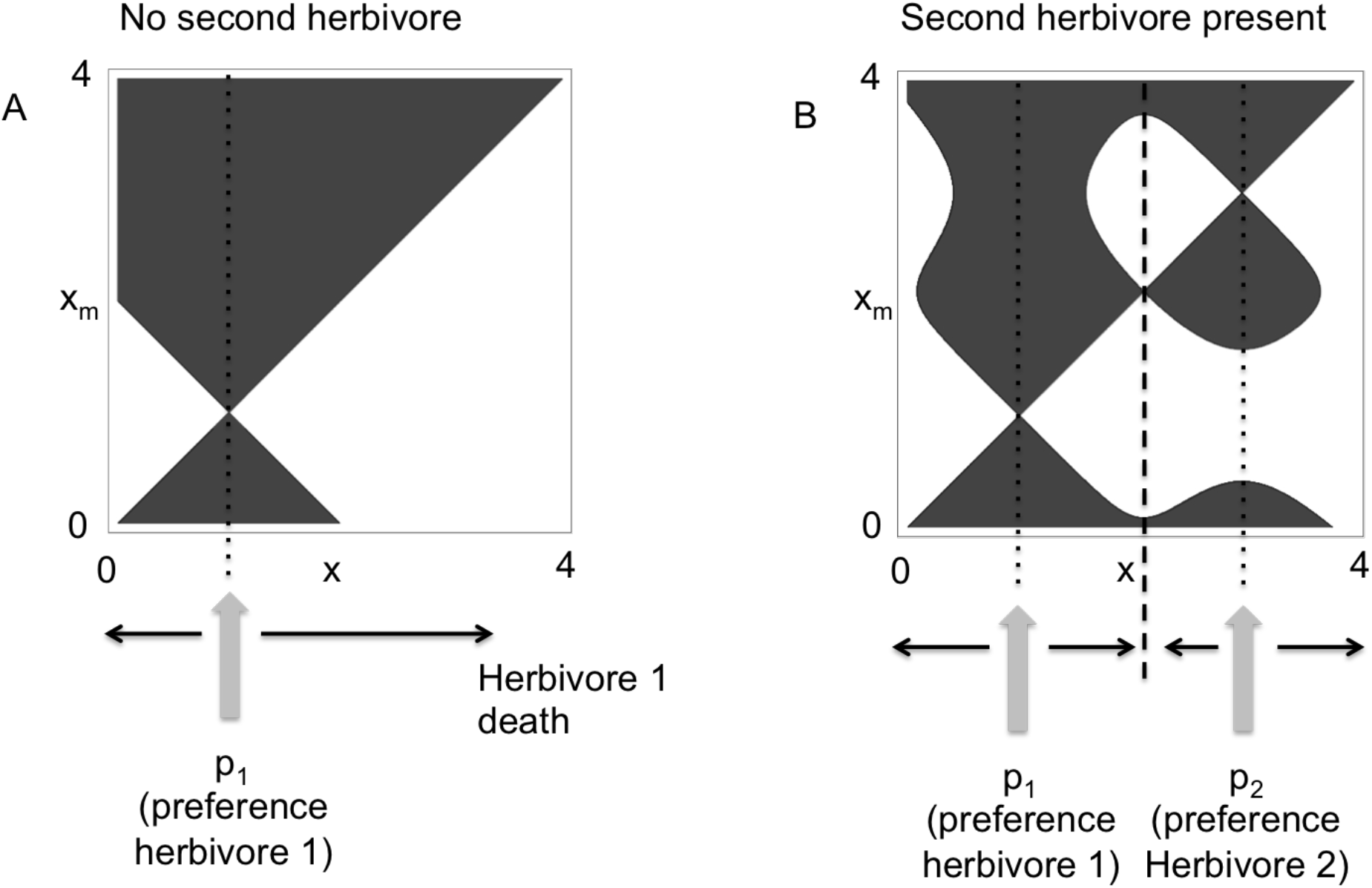
Effect of herbivore diversity on the evolution of qualitative defenses. Here, herbivore populations are considered constant (eg, herbivore populations vary on a much longer timescale). Thick grey arrows show the herbivore preferences. Black thin arrows show the direction of evolutionary dynamics of qualitative defenses. Dotted lines show the positions of the repellers and dashed line the position of the CSS. A) No second herbivore *(H_2_=0)*. Plants evolve away from preference *p_1_*, decreasing the herbivore 1 feeding rate eventually threatening its maintenance. B) The second herbivore is present *(H_2_=0.05)*. Due to its preference *p_2_*, evolution of the plant may settle between the two preferences, facilitating the coexistence of the two herbivores.

